# Defining cell type-specific diurnal gene expression in the mouse retina

**DOI:** 10.64898/2026.10.29.762877

**Authors:** Makayla N Marlin, Gaoya Meng, Shaherah Alqahtani, Patrick C. Kerstein, Vikki M. Weake

## Abstract

The mammalian retina exhibits daily oscillations in gene expression that contribute to retinal physiology and visual function. However, the extent to which these temporal patterns differ among retinal cell types remains incompletely understood. We performed single-nucleus RNA sequencing in the mouse retina over the day and night, identifying rhythmic genes in the six major retinal cell types. Distinct diurnal gene expression patterns across photoreceptors, interneurons, retinal ganglion cells, and Müller glia correspond to their diverse functions across the light cycle. Even core circadian clock genes exhibited differences in paralog expression and rhythmicity across the retina. In contrast, dopamine signaling via the DRD4 receptor is highly rhythmic across most retinal cell types, suggesting dopamine signaling may be a key rhythmicity cue in the retina under diurnal conditions. This dataset highlights the complexity of daily transcriptional regulation in the retina and provides a resource to explore cell-type-specific diurnal rhythms.

## Introduction

The vertebrate retina responds to light both for image-forming vision and for non-image forming visual processes, such as the pupillary light response and circadian photoentrainment ^1,2^. In mammals, light sensed by the retina provides a critical zeitgeber, or time-giver, to entrain daily biological rhythms – termed circadian rhythms ^3^. Visual signaling involves distinct cell types within the mouse retina, a highly organized and stratified tissue that contains photoreceptor cells (rods and cones), interneurons (amacrine and bipolar cells), horizontal cells, retinal ganglion cells (RGCs), and Müller glia ^4^. While the rod and cone photoreceptors detect low and bright light, respectively, for image-forming vision, a subset of RGCs, intrinsically photosensitive retinal ganglion cells (ipRGCs), detect ambient light to regulate circadian photoentrainment ^5^. Although light received by ipRGCs is critical to synchronize the central clock in the suprachiasmatic nucleus (SCN) ^6^, the retina itself has its own molecular clocks that independently keep time ^7^. Indeed, many physiological processes in the retina are under circadian control including photoreceptor disc shedding, release of melatonin and dopamine, and electrical responses to light ^8^. Despite the highly rhythmic nature of the retina, the rhythms present within cell types in this complex tissue are unknown.

On a molecular basis, biological rhythms are generated by a transcription-translation feedback loop, the circadian clock, that regulates gene expression in collaboration with zeitgebers such as light and dietary intake ^9^. The transcription factors CLOCK and BMAL1 (encoded by *Arntl*) act as positive regulators that drive transcription of clock output genes that encode their own negative regulators, PERIOD and CRYPTOCHROME family proteins ^7^. A second feedback loop involves the CLOCK/BMAL1 targets *Rev-Erbα* and *Rorα* that encode competitive regulators of *Arntl1* itself ^7^. The circadian clock also contributes to rhythmicity because microarray analysis of bulk mouse eye tissue reveals that approximately 60% of rhythmic transcripts require BMAL1 for rhythmicity even under normal light-dark cycles ^10^. Moreover, disrupting biological rhythms via genetic manipulation of the circadian clock negatively affects eye and retina homeostasis. For example, global *Bmal1* and *Clock* mouse mutants exhibit age-associated phenotypes that affect the eye such as cataracts ^11,12^. In addition, retina-specific disruption of the circadian clock leads to progressive photoreceptor degeneration in both mice (cones) and in *Drosophila* ^13–16^. Disrupting biological rhythms may also exacerbate inherited retinal dystrophies because rod-specific *Bmal1* knockout increases degeneration in mouse models of retinitis pigmentosa ^17^. The circadian clock also plays a crucial role in retinal development where *Bmal1* contributes to neurogenesis timing, cone identity, and vascular growth, leading to altered retinal structure and function ^18–20^. Although the circadian clock is essential to maintain biological rhythms in the absence of zeitgebers, rhythmic gene expression in the retina is strongly influenced by light ^10^. Light itself plays a key role in retinal development even before the conventional visual system has fully matured: the fetal mouse eye responds directly to light through melanopsin-expressing ipRGCs, influencing neuronal number, hypoxia related signaling, and vascular development ^21^. Despite the key role for the circadian clock in the eye, there are some discrepancies between reports describing the cell type-specific expression pattern of core clock genes in the retina ^22,23^, while the role of the circadian clock within specific cells in the retina remains unclear.

Consistent with a critical role for biological rhythms in maintaining ocular health, bulk transcriptome analysis reveals widespread rhythmic gene expression in the eye. Under light-dark cycles, approximately 2000 genes show rhythmic gene expression in the mouse eye and retina ^10,24^. Likewise in primates, 758 genes are rhythmically expressed in the baboon retina across the day and night ^25^. Notably, retina gene expression rhythms are strongly altered by aging in mice and *Drosophila* ^15,26^, as well as in diabetic mouse models ^27,28^. Recent advances in single cell sequencing technology have enabled transcriptome characterization of the different cell types within the mammalian retina, identifying distinct gene expression profiles corresponding to their function in visual signaling ^29^. Additionally, different retinal cell types exhibit distinct transcriptional changes during aging or in disease ^28,30–32^. Here, we used single nucleus RNA-seq (snRNA-seq) to profile the gene expression rhythms of the adult mouse retina across the light-dark cycle. We found that cells in the retina exhibit highly distinct rhythms in gene expression that coordinate with both structure and function. These data provide a unique resource to interrogate the cell type-specific diurnal rhythms that maintain retinal homeostasis and health in the adult retinas.

## Results

### SnRNA-seq of the mouse retina across the diurnal cycle

Bulk transcriptome analysis of the mouse eye indicates that while many genes are rhythmic under the light-dark cycle, far fewer genes remain rhythmic under constant darkness ^10^. Thus, we profiled rhythmically expressed genes in the mouse retina across the day under diurnal. We acclimated adult C57BL/6 mice under 12h light:12h dark (diurnal) conditions for 6 weeks and then collected retinas every 4 hours over a 24-hour period (Fig. 1A). Retinas were collected from two male and two female 4-month-old mice at each of the indicated zeitgeber times (ZT), beginning at ZT2, 2 hours after light onset. We performed snRNA-seq and identified six major retinal cell types across all integrated samples; rods, cones, bipolar cells, amacrine cells, RGCs, and Müller glia (Fig. 1B, C, Table S1). The retinal pigment epithelium (RPE) was not included in our retina dissections and was therefore underrepresented in our snRNA-seq data together with other low abundance cell populations, including horizontal cells. Although we identified subtype-specific markers such as *Opn4* (encoding melanopsin) for the ipRGCs and *tyrosine hydroxylase* (*TH*) for dopaminergic amacrine cells, these sub-populations lacked a sufficient number of cells across all samples to power rhythmicity analysis (Supp. Fig. 1).

**Figure 1.**
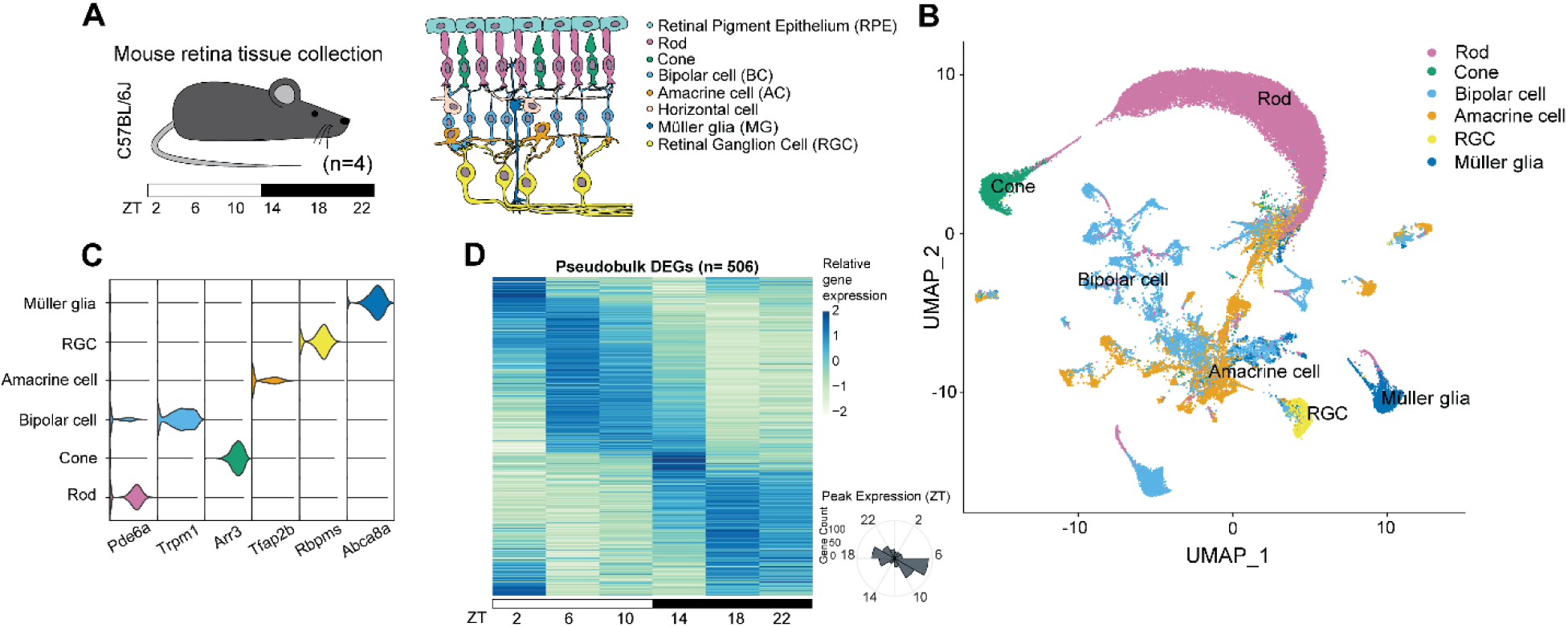
Major retinal cell types across the day identified by snRNA-seq. (A) Single-nucleus RNA sequencing (snRNA-seq) was performed on mouse retinas collected every 4 hours across a 24-hour light-dark cycle (ZT2, 6, 10, 14, 18, and 22; *n* = 4 biological replicates per time point, 2 females and 2 males). (B) UMAP embedding of snRNA-seq data from the retina where each point represents an individual cell, colored by cell type. (C) Violin plots showing expression of representative marker genes across annotated retinal cell types. (D) Pseudobulk heatmap of differentially expressed genes (DEGs) defined as rhythmic across the day (FDR < 0.05). DEGs were identified using aggregate expression counts from biological replicates at each ZT. Z score of variance stabilized expression values is shown. Radial histogram showing the phase of gene expression.

We first identified genes that exhibited differential expression across the day in the entire retina snRNA-seq dataset by combining all nuclei from each biological replicate regardless of cell type. Aggregate pseudobulk analysis of the retina revealed 506 genes that were differentially expressed across the day (Fig. 1D, Table S2). Most of these genes exhibited phases in the middle of the day (ZT6) or at night (ZT18). We re-analyzed bulk retina RNA-seq data from young C57BL/6 mice ^24^ and found that 158 genes were differentially expressed across the day in both datasets (Table S3; Supp. Fig. 2A and B). Since our data are based on nuclear RNA, these may more closely reflect nascent transcription compared with this previous bulk RNA-seq analysis that measures steady-state mRNA levels and can often be dominated in the retina by the signal from rod photoreceptors.

### Retinal cell types display distinct diurnal gene expression profiles

We next identified genes that were differentially expressed across the day in each of the six major cell types profiled, by aggregating samples from each ZT based on cell type, hereafter referred to as *rhythmic genes*. Rods contained the highest number of rhythmic genes (1259 genes) that exhibited dominant oscillating expression patterns, or phases, at ZT6 and ZT18 (Fig. 2A, Table S4), similar to our pseudobulk analysis of the entire retina. 325 of the rhythmic genes from rods were also identified as rhythmic in the bulk RNA-seq analysis of mouse retina ^24^ (Supp. Fig. 1C), consistent with the idea that rods heavily influence detected rhythmicity of genes in bulk retina data. Bipolar cells were the second most rhythmic cell type with 568 rhythmic genes, followed by cones (281 genes), amacrine cells (199 genes), Müller glia (108 genes), and RGCs (38 genes). These rhythmic patterns of gene expression were also apparent in both male and female mice across individual samples (Supp. Fig. 3). The lower number of rhythmic genes in RGCs likely represents diminished power due to the smaller number of RGCs in our dataset compared with more abundant cell populations. Comparison of the phase of rhythmic genes across these cell types revealed distinct patterns of rhythmic gene expression within the retina (Fig. 2A). Rods and cones followed a very similar pattern, with a strong peak in rhythmic gene expression at night (ZT18) and a smaller peak during the middle of the day (ZT6). In contrast, bipolar cells and amacrine cells had a peak in rhythmic gene expression predominantly in the early morning at ZT2, while Müller glia and RGCs follow a more distributed phasic pattern throughout the day (Fig. 2A).

**Figure 2.**
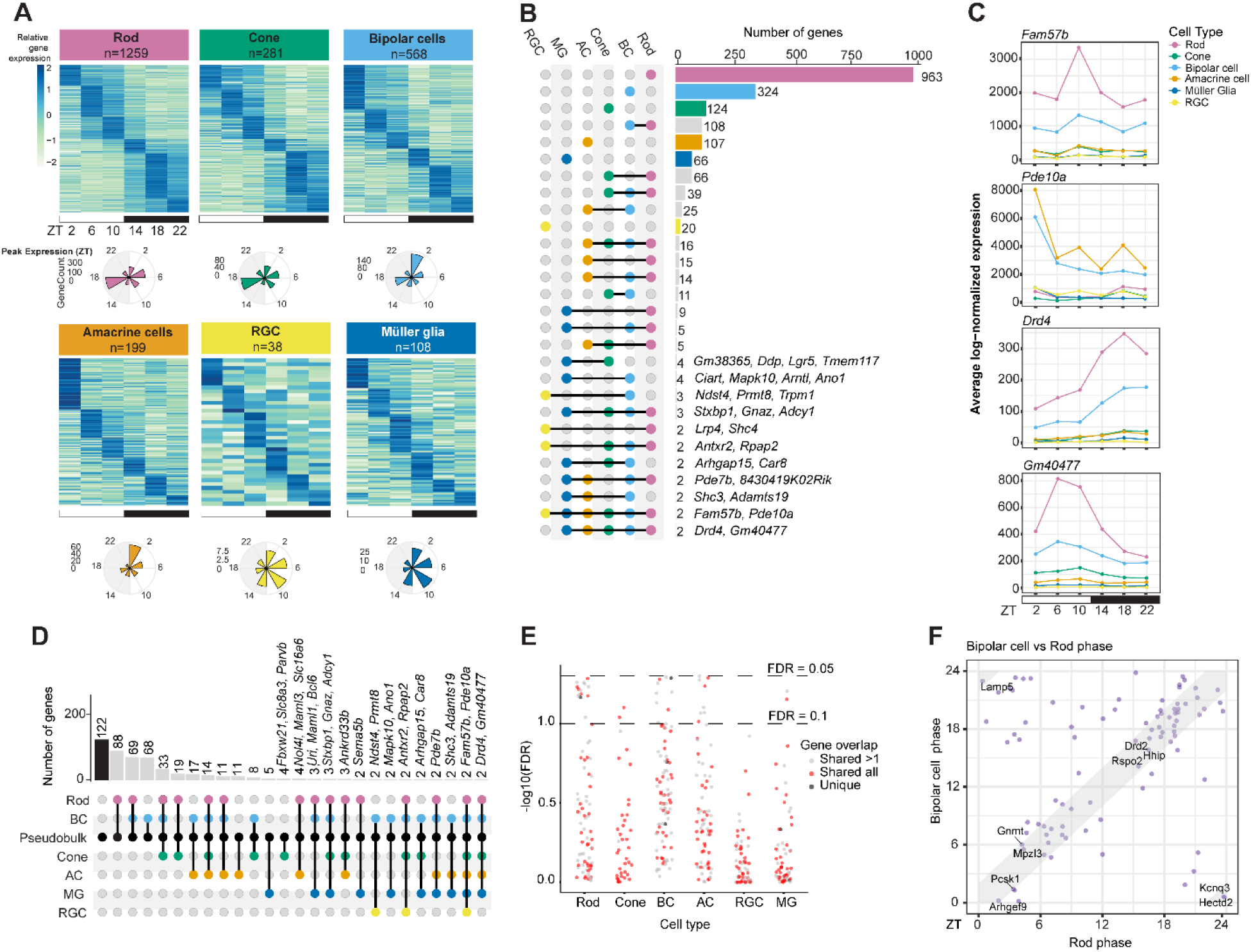
Cell type-specific diurnal transcriptional programs across the retina. (A) Heatmaps of differentially expressed genes (DEGs) defined as rhythmic across the day (FDR < 0.05). DEGs were identified using aggregate expression counts from biological replicates at each ZT. Z score of variance stabilized expression values is shown. Radial histogram showing the phase of gene expression. (B) UpSet plot indicating cell type-specific unique and overlapping rhythmic genes for groups ≥ 2 genes. (C) Line plots illustrating normalized expression of overlapping rhythmic genes in retinal cell types across the light-dark cycle. (D) UpSet plot showing unique and overlapping rhythmic genes in the pseudobulk analysis of the whole retina versus individual retinal cell types. (E) Dot plot showing FDR values for rhythmicity in each retinal cell type of the 122 rhythmic genes unique to the pseudobulk analysis of the entire retina. Individual genes (dots) are colored red if expressed in all cell types, black if uniquely expressed in a single cell type, and grey if expressed in more than one cell type. Dashed horizontal lines indicate FDR thresholds of 0.05 and 0.10. (F) Pairwise scatterplot compares expression phase in bipolar cells versus rods for the 91/122 rhythmic genes expressed in these cell types. Genes falling within the gray-shaded diagonal region exhibit similar phases ±2 hours.

Unexpectedly, relatively few genes were rhythmic in multiple cell types and only two genes were rhythmic in all six cell types: *Fam57b* and *Phosphodiesterase 10A (Pde10a)* (Fig. 2B). *Fam57b*, also referred to as TLCD3B, encodes the most highly expressed ceramide synthase in the human retina and has been associated with human recessive retinal dystrophy ^33^. *Fam57b* expression peaks at ZT10 with the highest expression in rods and bipolar cells (Fig. 2C). *Pde10a* encodes a phosphodiesterase that hydrolyzes cyclic AMP (cAMP) and cyclic GMP (cGMP) to regulate dopaminergic and cAMP/protein kinase A (PKA) signaling cascades ^34^. *Pde10a* transcript and protein levels are highly rhythmic in rat retina, with these rhythms sensitive to altered light and dark cycles ^35^. In our data, amacrine and bipolar cells had the highest expression of *Pde10a* at ZT2 in the early morning (Fig. 2C). Two additional genes were rhythmic in every cell type except for RGCs: *Dopamine receptor d4* (*Drd4*) and *Gm40477*. *Drd4* encodes a G-protein coupled receptor that binds dopamine, a critical neurotransmitter for light signaling in the retina ^36^. DRD4 signaling suppresses light-sensitive cAMP in photoreceptors via type 1 adenylyl cyclase and, in coordination with D1 receptor signaling, contributes to light adaptation, contrast sensitivity, and the regulation of retinal circadian rhythms ^37,38^. *Drd4* peaked in expression during the night at ZT18 in rod and bipolar cells (Fig. 2C). *Gm40477* is a predicted long non-coding RNA that has been annotated as a *Fos* enhancer RNA ^39^, and peaked in expression at ZT6 in rod and bipolar cells (Fig. 2C).

In contrast to the relatively small number of genes with shared rhythmicity across multiple cell types in the retina, most rhythmic genes were cell-type specific or only shared between two functionally related cell types. For example, rods and bipolar cells had 963 and 324 unique rhythmic genes, respectively (Fig. 2B). A further 108 genes were rhythmic in both rods and bipolar cells, while 66 genes were rhythmic in rods and cones. These highly cell type-specific rhythms provide an explanation for the substantially larger number of unique rhythmic genes identified collectively within the individual retinal cell types (1604 genes) compared with the aggregate (pseudobulk) analysis of the same retina dataset. Indeed, most of the 506 rhythmic genes identified in the pseudobulk retina data represent genes that were rhythmic in either rods or bipolar cells, the two most abundant cell types in our dataset (Fig. 2D). Interestingly, 122 genes were classified as rhythmic only in the pseudobulk analysis, but not in the individual cell types (Fig. 2D). Many of these 122 pseudobulk-specific rhythmic genes were expressed in multiple retinal cell types, with 9% of these genes just falling short of the significance threshold in any one cell type (Fig. 2E). Since a number of these genes (n=91) also exhibit similar phases (∼45% ± 2 hours) in more than one cell type, such as in rods and bipolar cells (Fig. 2F), their shared phase across several cell types may increase statistical power for detecting rhythmicity. This comparison also highlights the fact that gene rhythms may be masked both by sub-type heterogeneity and limitations in detecting differential expression within some less abundant populations.

### Cell type-specific rhythms in the retina temporally separate visual processes

We performed Gene Ontology (GO) term enrichment analysis by cell type across the day, showing that rhythmic genes in the retina were associated with diverse biological processes at distinct times of the day (Fig. 3, Table S5). There are two types of photoreceptors: rods and cones. Rods detect light for scotopic or low light vision; in contrast, cones detect light and color for photopic vision ^40^. During the light phase, rhythmic genes in rods were associated with *ribosome biogenesis* and *DNA damage*, potentially reflecting increased biosynthetic and cellular maintenance demands associated with photoreceptor outer segment renewal, and protection from light- and metabolism-associated oxidative stress ^41,42^. During the early evening, rods expressed genes that *respond to oxygen*, consistent with the increase in oxygen consumption during the night to meet the demands of ATP-dependent ion transport ^42^. Later in the night, rods express genes involved in *metabolic processing of purine nucleotides and glycoproteins*, consistent with increased metabolism in the dark phase, which coincides with mice wakefulness ^43,44^. In cones, rhythmic genes expressed during the light phase are involved in *sensory perception of light stimulus* and *protein localization to the cell membrane*. Cone phototransduction is active during photopic day-time conditions, and protein sorting and trafficking to the membrane is essential for maintaining the phototransduction machinery in these highly polarized photoreceptor cells ^45^. Cones also express rhythmic genes involved in *synapse assembly* at the end of the night followed by *axon regeneration* genes early in the morning, potentially contributing to the dynamic remodeling of activity-dependent cone circuitry during the light phase ^46^. In the early evening, cones express genes involved in *ketone metabolism* – potentially as part of the shift from metabolizing glucose ^47^.

**Figure 3.**
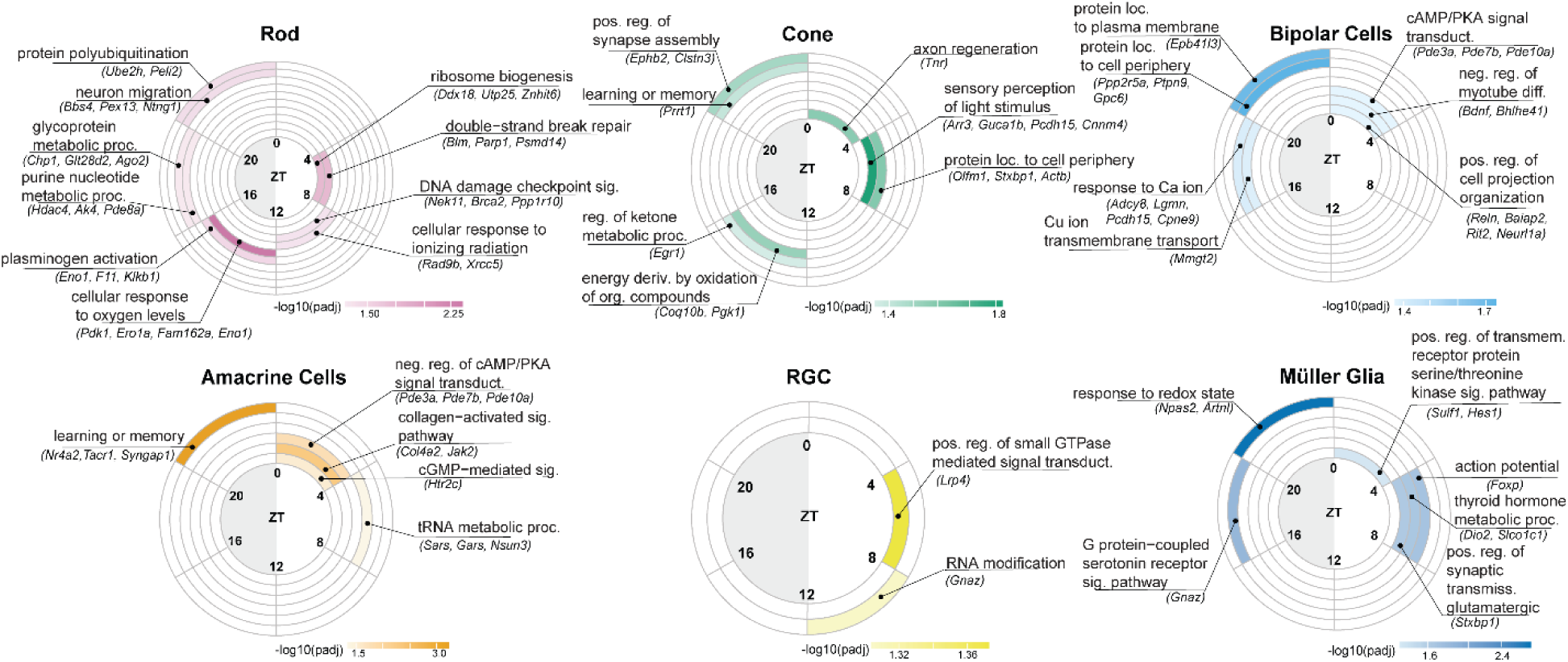
Cell type-specific temporal rhythms in retinal visual processes. Circular plots summarize the top three gene ontology (GO) biological processes associated with genes peaking at different phase windows within each retinal cell type. Colored by -log10(padjust) values.

Within our dataset, we analyzed two types of interneurons: bipolar cells and amacrine cells. Bipolar cells are excitatory interneurons that connect the outer retina (rods and cones) to the inner retina and form two subclasses that depolarize (ON) or hyperpolarize (OFF) to increments in light intensity ^40^. These subtypes are combined in our data showing rhythmic expression in the morning of genes involved in *cAMP/PKA signal transduction*, which modulates bipolar-cell responsiveness to changing photoreceptor input during the transition to daytime vision ^48^. *Cell-projection organization* genes are also expressed in bipolar cells in the morning, including regulators of the structural and cytoskeletal machinery that maintains bipolar-cell dendrites and axon terminals ^49^. The night-time expression of genes involved in the *response to calcium ions* may allow bipolar cells to respond to the increased activation of L-type voltage gated calcium channels at night ^50^. Amacrine cells modulate communication between bipolar cells and RGCs, providing inhibitory and neuromodulatory processing that enables temporal, spatial, and motion related vision ^40^. In the light, amacrine cells, like bipolar cells, are enriched for genes involved in *cAMP/PKA signal transduction* and *cGMP-mediated signaling*. The enrichment of these pathways may be involved in light stimulation of dopamine signaling in amacrine cells ^51^, increasing intracellular cAMP and PKA-mediated phosphorylation that can be modulated by cGMP and phosphodiesterase activity ^52^. During the night, amacrine cells express genes involved in *learning and memory* processes that are associated with activity-dependent synaptic plasticity. This may reflect the dynamic modulation of amacrine cell function by retinal activity and illumination.

RGCs are the primary output neurons of the retina that integrate signals from bipolar and amacrine cells and send action potentials through the optic nerve to visual centers in the brain ^40^. In the light phase, RGCs express *regulators of small GTPase mediated signal transduction*, driven by LRP4, which itself is a small GTPase, but can influence signaling indirectly by transcription factors whose targets include GTPase regulators ^53^. While the GO term enrichment for RGCs was limited by the relatively small number of rhythmic genes identified in this cell type there were 20 uniquely rhythmic genes in RGCs (Fig. 2B). These include cell adhesion molecules *Cntnap5a* and *Epha3* that peaked at ZT2 and belong to gene families previously implicated in synapse formation and remodeling ^54,55^.

Müller glia span the entire thickness of the retina, providing structural, metabolic and extracellular homeostatic support to retinal neurons ^56^. In the light phase, Müller glia expressed genes involved in *kinase signaling*, *action potential*, and *glutamatergic synaptic transmission*, which may be associated with the circadian regulation of glutamate signaling in the retina and modulation of homeostasis. During the dark phase, Müller glia expressed genes involved in the *response to redox state*, potentially as part of their role as major regulators of the retinal redox environment ^57^. Müller glia have transcriptional programs that are temporally regulated by redox status, and oxidative-stress responses oscillate across the light cycles ^58^. Together, these results demonstrate that diurnal gene expression in the retina is highly cell-type specific, with distinct temporal programs coordinating the metabolic, signaling, and homeostatic functions of individual retinal cell types across the light-dark cycle.

### A subset of rhythmic genes exhibit distinct phases across retinal cell types

Although many genes were uniquely rhythmic in one retinal cell type, some genes showed overlapping rhythmicity in two or more cell types, such as the 108 genes that were rhythmic in both rods and bipolar cells (Fig. 2B). Some of these shared rhythmic genes had similar phases in both cell types, such as the genes involved in *cAMP/PKA signal transduction* that were expressed in the morning in both bipolar and amacrine cells (Fig. 3). However, we wondered if other rhythmic genes might exhibit distinct phases in different retinal cell types. Indeed, there were varying degrees of phase consistency among shared rhythmic genes between cell types (Fig. 4A). Although some shared rhythmic genes exhibited highly concordant peak expression times between two cell types, a subset of genes were expressed at distinct circadian phases in different cells. These distinct phases were most apparent between rods and amacrine cells, and rods and bipolar cells where many genes show a shift in phase 12h later in rods compared to the other two cell types. Of note are some of the circadian genes, such as *Cry1* and *Per2*, that have phases near ZT18 in rods, but around ZT6 in amacrine cells. Intriguingly, one of the genes that is rhythmic in all six retinal cell types, *Pde10,* had distinct phases between rods and most other cells, peaking near ZT18 in rods compared to ZT6 for amacrine and bipolar cells, and Müller glia.

**Figure 4.**
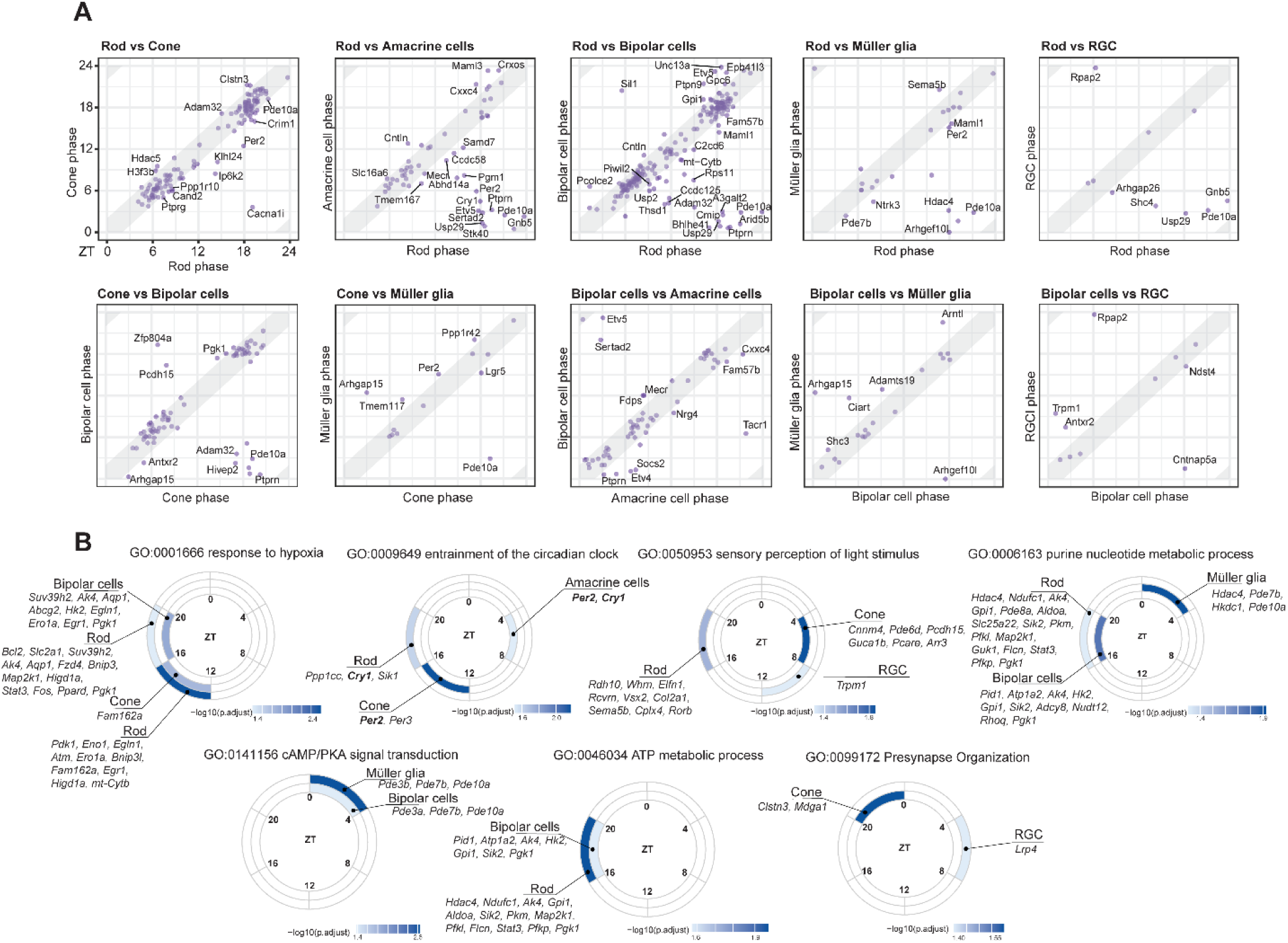
Peak expression timing reveals coordinated functional programs. (A) Pairwise scatterplots comparing the phase of shared rhythmic genes between retinal cell types. For each shared gene, the peak expression phase in one cell type was plotted against its corresponding phase in the second cell type. Genes falling within the gray-shaded diagonal region exhibit similar phases of expression (within ±2 hours) between the two cell types, whereas genes outside this region display divergent phases. (B) GO terms shared across cell types are displayed with enrichment of rhythmic genes shown across the diurnal day. Cell types are displayed at time points where the corresponding GO term was identified.

These differences in gene-specific rhythms correspond with distinct timing of the same biological process across the retina. For example, *response to hypoxia* genes are expressed in cone and rods from ZT12-16, but in bipolar cells from ZT16-20 (Fig. 4B). This suggests that there may be a lag in oxygen demands between rods and cones, and their synaptic partners, bipolar cells, during the early stages of the night. Further, *sensory perception of light stimulus* occurs in cones during the day at ZT4-8, but in rods at ZT16-20 during the night. This likely reflects the time of day in which these photoreceptors are more responsive to different light levels ^59^. Thus, differences in the temporal regulation of gene expression across the day between cell types likely reflect functional differences related to the light response.

### Circadian clock genes exhibit distinct diurnal patterns across retinal cell types

Unexpectedly, we observed distinct phases of circadian clock genes (*GO: 0009649*) in rods, cones, and amacrine cells (Fig. 4B), suggesting that the circadian clock may be differentially regulated across the retina. In mammals, *Clock/Npas2* and BMAL1 (encoded by *Arntl*) activate transcription of *Cry1/2, Per1/2/3, Nrd1/2* (encoding REV-ERB), and *Dbp* (Fig. 5A). DBP in turn activates transcription of *Rora/b/c* (encoding ROR), which competitively regulates *Arntl* transcription with REV-ERB ^60^. Our snRNA-seq data suggests that most retinal cell types express *Clock* and *Arntl* (Fig. 5B), consistent with published single cell RNA-seq data from the adult mouse retina ^29^ (Fig. 5C). However, there are some differences in relative expression of clock output genes between the cell types with most retinal cells exhibiting higher expression of *Per3* compared with *Per1* and *Per2*. While *Per1/2* play a critical role in the circadian clock in the SCN and are required for behavioral rhythms in mice, *Per3* often plays a more tissue-specific modulatory role in fine-tuning rhythms ^61^. Similarly, *Cry2* expression was higher in many of the retinal cell types except for rods, where *Cry1* expression was higher. Notably, even though many of the circadian clock genes are expressed throughout the retina, not all exhibit rhythmicity in each cell type. Each cell type except for RGCs showed rhythmic expression of at least two core circadian clock genes, but not necessarily the same genes. For example, *Per2* was rhythmic in rods, cones, amacrine cells, and Müller glia, while *Per1* was rhythmic in bipolar cells. Although more highly expressed across the retina, *Per3* was only rhythmic in cones. Similarly, while *Cry2* expression was higher in many retinal cell types, only *Cry1* was rhythmic in rods and amacrine cells. Moreover, *Arntl* was only rhythmic in bipolar cells and Müller glia. The lack of rhythmicity of core clock genes in RGCs potentially reflects the low abundance of this cell population in our dataset. We obtained similar results even when using a second rhythmicity analysis approach, *dryR* ^62^ (Table S6). The differences in rhythmicity of core clock genes across the retina implies there are functional differences in the nature of the transcription-translation feedback loop in each cell type.

**Figure 5.**
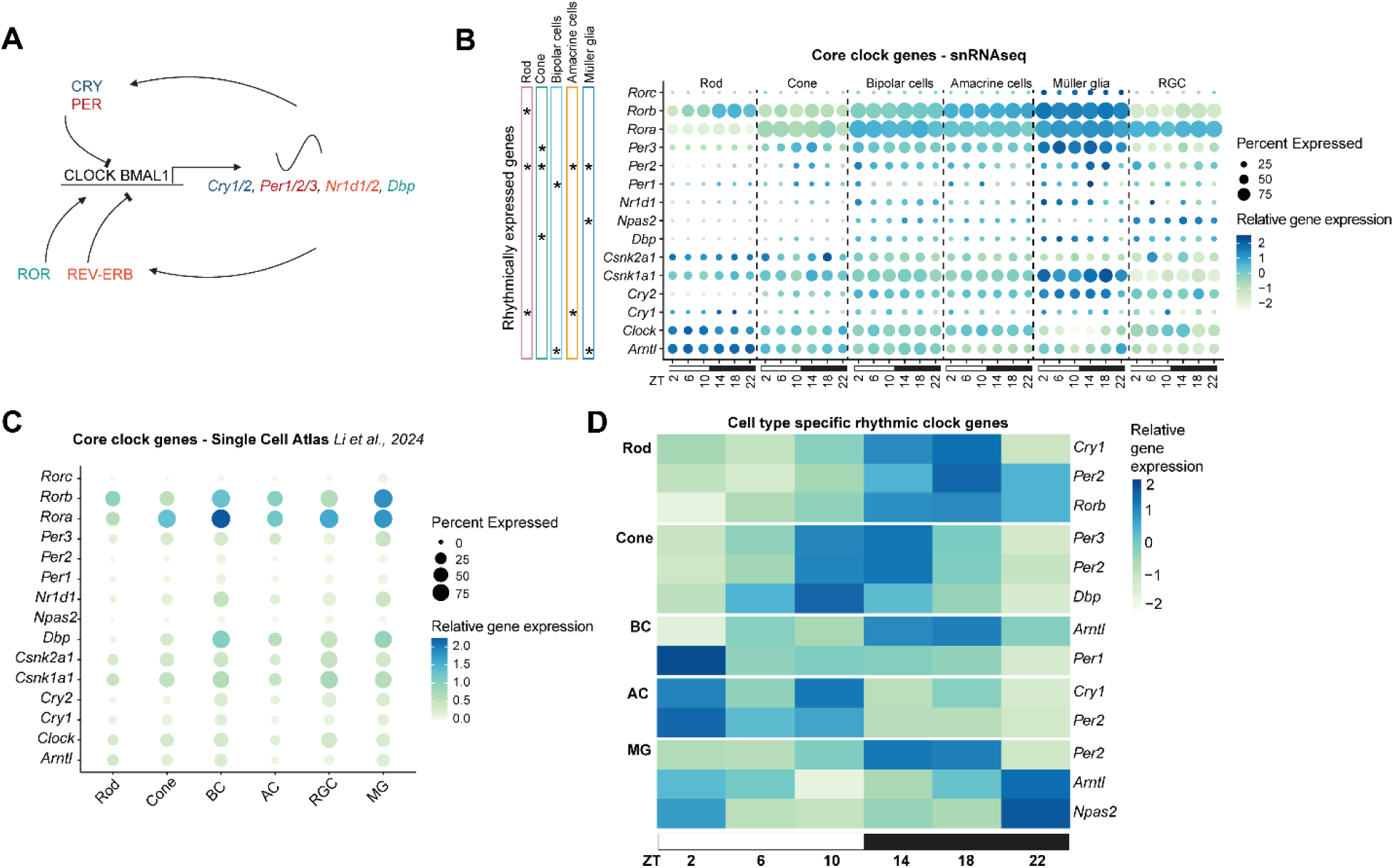
Distinct rhythms of core clock gene expression across the retina. (A) Schematic for core molecular clock. CLOCK–BMAL1 activates *Per* and *Cry* transcription through E-box motifs, while PER/CRY proteins inhibit CLOCK–BMAL1 through negative feedback. ROR and REV-ERB regulate *Bmal1* transcription, promoting and repressing its expression, respectively. (B) Dot plots showing the relative expression of core clock genes across retinal cell types at each ZT. Dot size represents the percentage of nuclei within each cell type expressing the gene, while dot color indicates the average normalized expression level. Rhythmic genes (FDR < 0.05) are denoted by an asterisk. (C) Dot plot of core clock gene expression in the six major retinal cell types based on single cell RNA-seq data from the single cell atlas ^29^. (D) Heatmap (Z-score) of rhythmically expressed core clock genes across the diurnal day.

Intriguingly, there seemed to be some differences in the phase of core clock output genes between closely related cell types in the retina. For example, *Cry1*, *Per2,* and *Rorb* peak at ZT18 in rods, while *Per2, Per3,* and *Dbp* peak 6 to 8h earlier in cones, with *Per2* in Müller glia exhibiting an intermediate phase at ZT15 (Fig. 5D, Table S4). These same core clock genes peak much earlier in the morning in amacrine cells, with *Cry1* and *Per2* peaking at ZT4-6 (Table S4). The phase of *Arntl*, the positive arm of the circadian oscillator, is typically different to that of clock output genes, and in Müller glia, *Arntl* and *Npas2* peak at ZT22, whereas in bipolar cells *Arntl* peaks at ZT18 ^63^. It remains unclear whether these differences in the phase of nuclear RNA expression reflect corresponding differences in clock function or downstream physiological activity. Nevertheless, these findings suggest that while the core molecular clock machinery is conserved across retinal populations, its rhythmic transcription regulation may be tailored to the functional requirements of individual cell types.

### Light-responsive dopamine signaling is diurnally regulated

Beyond the core circadian clock, the retina responds to the changing light cycle via neurotransmitters that coordinate neuronal function across the day ^3^. Dopamine is a key neuromodulator that responds to light and is released by dopaminergic amacrine cells in the inner plexiform layer to modulate chemical and electrical signaling across the light-dark cycle ^64^ (Fig. 6A). Retinal dopamine release is higher during the day and decreases at night and is regulated by the retinal circadian clock and light exposure, providing a mechanism for coordinating daily changes in retinal function ^59^. Dopamine acts through distinct receptor subtypes to modulate intracellular signaling. In photoreceptors, DRD4 functional activation inhibits adenylyl cyclase, reducing cAMP levels and PKA activity to help regulate light adaptation ^65^. Rhythmic dopamine signaling appears to be maintained in light-dark conditions even in retinas lacking melatonin, suggesting that local regulation of dopamine signaling may contribute to its daily rhythm ^66^. Indeed, *Drd4* is one of the most rhythmic genes across all cell types in the retina except for RGCs. Consistent with this, microarray analysis of the mouse eye revealed highly rhythmic *Drd4* transcript levels under light:dark conditions that persisted in the *Bmal1* mutant ^10^. Similarly, qPCR analysis of *Drd4* transcript levels in bulk retina tissue showed rhythmic expression under the light-dark cycle, with significant, albeit dampened, rhythm during constant darkness ^67^. Notably, whereas DRD4 functional activation inhibits adenylyl cyclase and reduces cAMP production, PDE10A hydrolyzes cyclic nucleotides including cAMP and cGMP ^35^. Interestingly, *Pde10a* is one of the two rhythmic genes that are expressed in all cell types and may contribute to the rhythmic regulation of dopamine signaling through its role in cyclic nucleotide signaling downstream of DRD4 activation. In most retinal cell types in our data, *Drd4* peaked around ZT18, whereas *Pde10a* peaked around ZT2 in all but the photoreceptors where it peaked at ZT22 (Table S4).

**Figure 6.**
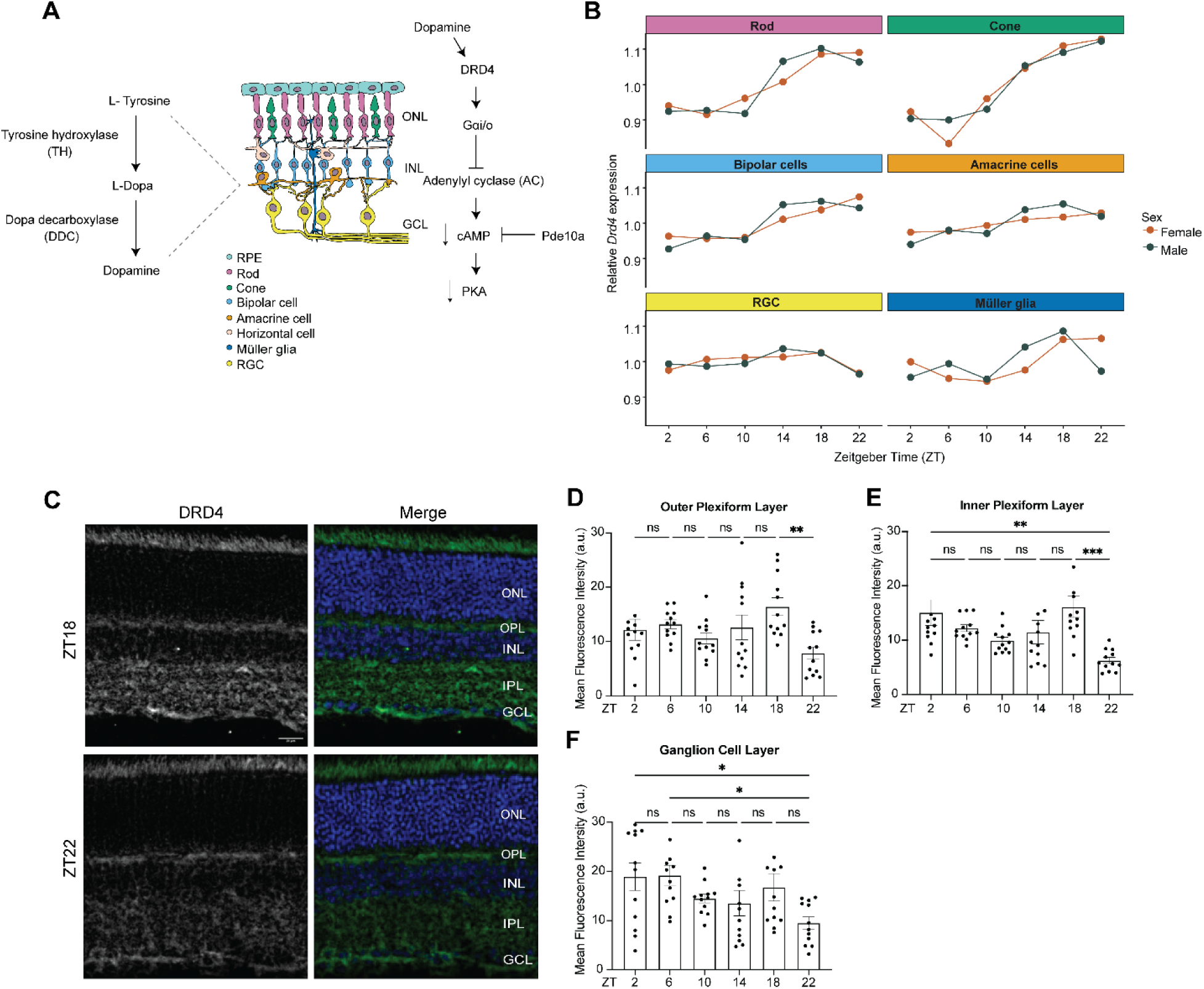
The dopamine receptor Drd4 involved in light-responsive dopamine signaling is diurnally expressed. (A) Schematic illustrating the pathways involved in dopamine production and release by dopaminergic amacrine cells that subsequently activates dopamine D4 receptors (DRD4) in the retina. (B) Relative *Drd4* expression across the diurnal cycle in each retinal cell type, shown separately for male and female mice, with expression averaged across biological replicates within each sex. (C) Representative images showing DRD4 protein expression in the mouse retina with the outer nuclear layer (ONL), outer plexiform layer (OPL), inner nuclear layer (INL), inner plexiform layer (IPL), and ganglion cell layer (GCL) indicated. DRD4 immunofluorescence (green) is shown relative to DAPI (blue). (D-F) Quantification of DRD4 mean fluorescence intensity across ZT in the OPL (D), IPL (E), and GCL (F) (n=12 measurements, n=2 mice). Mean fluorescence is plotted as a function of ZT for each retinal layer. *p<0.05, **p<0.01, ***p<0.005 represent statistically significant as evaluated by a One-way ANOVA with a multiple comparisons test. Ns, not significant. Scale bar, 20 µm in (C).

We observe rhythmic expression of *Drd4* across most cell types in the retina except RGCs, with patterns in male and female mice showing only subtle differences (Fig. 6B). In our nuclear RNA, *Drd4* has a phase of ZT18 in most cell types, while the bulk retina RNA-seq analysis ^24^ shows a phase of ZT19 (Table S3). We examined DRD4 protein levels in the retina across the day by immunostaining retinas for each time point (Fig. 6C-F). DRD4 showed strong signals in the outer and inner plexiform layers, and ganglion cell layers (Fig. 6C). Therefore, we quantified the immunofluorescence intensity of DRD4 within these layers across six timepoints of the light-dark cycle in both male and female mice (Fig. 6D-F). Like we observed in most cell types from our snRNA-seq analysis, we found that DRD4 immunofluorescence peaks at ZT18 in both the outer and inner plexiform layers (Fig. 6C-E). Interestingly, the peak of DRD4 protein from immunofluorescence in the ganglion cell layer peaked earlier at ZT2 instead of ZT18 (Fig 6F). However, since *Drd4* was not identified as rhythmic in RGCs based on our snRNA-seq data, it remains unclear if this difference in phase reflects distinct *Drd4* transcription rhythms between these retinal cell types. Despite this, our data suggest that dopamine signaling proteins are coordinately regulated across the day in many retinal cell types, highlighting a potential role for light-dependent dopamine release in these retinal rhythms.

## Discussion

Previous studies have established that the retina contains an intrinsic circadian clock and that light-responsive pathways influence retinal physiology and development. However, the extent to which circadian regulation is coordinated across individual retinal cell types has remained less clear. Here, using snRNA-seq across the 24-hour day, we identified extensive cell-type specific rhythmic gene expression in the mouse retina, revealing distinct temporal programs across photoreceptors, interneurons, RGCs, and Müller glia.

There have been some discrepancies in reports describing the expression of core clock components in the mouse retina, with variations in cell-type localization, expression, and rhythmicity ^8^. RT-qPCR analysis identified coordinated rhythmic expression of core clock genes (*Per1, Per2, Cry1, Cry2, Clock* and *Arntl/Bmal1*) in inner-retinal neurons but failed to detect all the core clock genes in photoreceptors ^23,68^. In contrast, immunohistochemistry analysis showed rhythmic expression of CLOCK, BMAL1, NPAS2, PER1/2 and CRY2 in cones, but very low and arhythmic expression in rods ^22^. Our cell-type-resolved analysis resolves some of these questions by showing that while many clock genes are broadly expressed across the retina, their rhythmicity is often limited to a subset of retinal cell types. These differences might reflect subtle variations on the circadian clock machinery between retinal cell types; for example, the three different *Per* paralogues are each rhythmic in distinct retinal cell types – with *Per3* rhythmic in cones, *Per1* rhythmic in bipolar cells, and *Per2* rhythmic in all cell types except for bipolar cells and RGCs. While *Per3* expression was previously reported to be arhythmic in bulk retina and in the outer photoreceptor layer ^69,70^, its rhythmicity in cones may have been masked by the more abundant signal from rods. Functionally, these variants in the circadian clock machinery likely contribute to retinal rhythms because individual deletion of *Per1, Cry1* or *Clock* weakened molecular rhythms in retinal explants, whereas deletion of *Per3* did not abolish, but rather shortened the period of the clock ^71^.

Although the core circadian clock plays a critical role in regulating rhythmic gene expression and maintaining retinal health, as shown by the impact of *Bmal1* mutations, daily rhythms in the retina are also coordinated by neuromodulatory pathways such as dopamine signaling ^10^. Dopamine levels, including its metabolites 3,4-dihydroxyphenylacetic acid (DOPAC) and homovanillic acid (HVA), exhibit daily rhythms in mice maintained under 12-hour light:12-hour dark cycles, although the persistence of these rhythms under constant darkness depends on strain-specific differences in melatonin production ^66^. The sensitivity of retinal dopamine receptors may allow these daily fluctuations to regulate retinal function. Dopamine d4 receptors (*Drd4*) have a higher affinity for dopamine than D1 receptors (*Drd1*), potentially allowing modest changes in dopamine availability to influence D4-mediated signaling, whereas stronger dopamine release associated with bright light can recruit lower-affinity D1 receptors ^59^. Thus, rhythmic dopamine availability may provide a mechanism for modulating photoreceptor signaling across the light-dark cycle, highlighting *Drd4* as a potential link between daily dopamine rhythms and circadian regulation of retinal function. Phosphodiesterase expression and activity may also contribute to the temporal regulation of cyclic-nucleotide signaling in the retina with differences in the phase of *Pde10a* between the photoreceptors and other retinal cell types. The rhythmic expression of these components may therefore reflect coordinated regulation of intracellular signaling pathways involved in daily retinal adaptation ^35^.

A second neuromodulator, melatonin, also acts as a key regulator of biological rhythms and has an opposing rhythm to dopamine – higher in the night relative to the day as it is synthesized in the pineal gland at night ^72^. Melatonin interacts through a family of G-protein coupled receptors (MT1 and MT2) that primarily couple inhibitory Gi proteins to suppress adenylyl cyclase activity, thereby reducing intracellular cAMP levels and downstream PKA signaling which can signal to pathways that influence retinal physiology ^72^. Because the C57BL/6 mice used in our study have impaired melatonin synthesis ^73^, there may be additional rhythmic genes regulated via this neuromodulator in the retina, particularly at night. However, similar overall numbers of rhythmic genes were identified in the eye of melatonin proficient mice versus the retina of C57BL/6 mice under light:dark conditions, despite differences in technical and analytical approaches (microarray versus RNA-seq) ^10,24^. Moreover, MT1 and MT2 melatonin receptor knock-out mice showed differences in clock gene expression rhythms primarily under constant darkness, while most rhythms persisted under light:dark cycles albeit with some changes in phase ^74^. In addition, melatonin had a stronger impact on clock gene rhythms in the inner retinal neurons, with little effect on photoreceptors ^74^. Despite this, the deficiencies in melatonin production in this mouse background represent a caveat of the current study.

Beyond providing insight into the mechanisms that contribute to cell type-specific diurnal rhythms in the mouse retina, this study provides a rich resource for investigating diurnal regulation of retinal function. However, several limitations should be considered when interpreting this dataset. First, our analysis is based on snRNA-seq, which captures nuclear RNA, including nascent and incompletely processed transcripts, in addition to mature RNA retained in the nucleus ^75^. Consequently, the rhythmic patterns observed here may not fully reflect steady-state mature mRNA abundance in the cytoplasm. Second, rhythmic transcript abundance does not necessarily correspond to rhythmic protein abundance or activity, as differences as translation, protein stability and turnover, and post-translational regulation can alter the timing and magnitude of downstream molecular responses ^76^.

The distinct rhythms exhibited by each cell type in the mouse retina raise important questions about how these rhythms change during aging and disease. During aging, losses in rhythmic gene expression have been observed in old mouse retinas ^26^. Likewise, in retinal disease, changes in rhythmic gene expression have been observed in mouse models of diabetic retinopathy ^27,28^. In these models, dopamine signaling may play a central role since dopaminergic amacrine cells that synthesize dopamine are susceptible to both dysfunction and cell death in the diabetic retina ^77,78^. Notably, our analyses demonstrated a rhythmic pattern among genes associated with dopamine signaling (*Drd4* and *Pde10a*) further highlighting the importance of studying rhythmic gene expression in models of retinal disease. These findings may extend beyond the retina, as disrupted dopamine signaling and circadian rhythms are both features of neurodegenerative disorders such as Parkinson’s disease ^79–81^. Defining cell-type-specific rhythms in the healthy retina provides a baseline for identifying cells and pathways that may lose rhythmic control in aging and disease ^13,25–27^.

## Materials and Methods

### Animals

The C57BL/J mouse strain was obtained from the Jackson Laboratory. All animal procedures were approved by the Purdue Animal Care and Use Committee (PACUC) at Purdue University and were conducted in accordance with the NIH Guide for Care and Use of Laboratory Animals. Mice were housed in standard 12-hour light: 12-hour dark (LD) conditions with standard laboratory rodent chow and water available *ad libitum*. To facilitate tissue collection during normal working hours, one cohort was maintained on a reversed LD cycle (lights on and off shifted by 12 h relative to the standard LD cycle). Animals were acclimated to their lighting schedules for 6 weeks before sample collection at about 4 months old. All tissues were collected at ZT defined as relative to lights-on for each housing room.

### Tissue collection and immunostaining

Animals (n=2 male, n=2 female) were euthanized using CO_2_ followed by cervical dislocation at ZTs 2, 6, 10, 14, 18, and 22. Collections during the dark phase were performed under dim red light. Eyes were rapidly enucleated after euthanasia and placed in Phosphate Buffered Saline (PBS). The cornea, lens and vitreous were removed and retinas were dissected and immediately flash frozen in liquid nitrogen and stored at −80°C for snRNA-seq. For immunohistochemistry, the corneas were removed, and the eye cups were placed in 4% EM-grade paraformaldehyde (PFA) for 30 mins at room temperature. The eyes were then washed 3 times in PBS for 10 mins. Next, Eyes were cryo-preserved by placing them in 15% sucrose for at least 2 hours at 4°C and then embedding them in a cryo-mold in Optimal Cutting Temperature media and flash frozen. Using a cryostat, 20 µm retinal sections were collected on a glass microscope slide (Fisher Superfrost Plus). For immunolabeling, retinal sections were washed in 1X PBS for 10 mins and then blocked with 2% normal donkey serum (NDS) with 0.2% Triton X-100 for 30 mins. Retinal sections were incubated with primary antibodies in NDS blocking buffer overnight at 4°C. A primary antibody against DRD4 (Invitrogen #PA5-104385) was used at a 1:200 dilution. Sections were then washed in NDS blocking solution and incubated with Alexa Fluor 488 Donkey anti-Rabbit secondary antibody (ThermoFisher #A-21206) for 2 hours at room temperature. Sections were washed three times in PBS for 10 mins with 4′,6-diamidino-2-phenylindole (DAPI) (1:5000) stain included in the first wash step. Finally, tissue was mounted in Fluoromount-G (Southern Biotech) and cover-slipped for imaging by Zeiss LSM900 Confocal Microscope.

### snRNA-seq

Retinas were homogenized in ice-cold nuclei isolation buffer using a glass Dounce homogenizer, and nuclei prepared using the Illumina Single Cell Nuclei Isolation kit (Catalog #20132795), as described by the manufacturer’s protocol. Nuclei were stained using Invitrogen ReadyCount stains (A49905) and quantified using Countess II (Invitrogen). Single-nuclei libraries were prepared in batches of 8 samples using the Illumina Single Cell 3’ RNA Prep T10 kit (Catalog #20132791, 20135691) according to the manufacturer’s protocol. Approximately 17,000 nuclei were used per sample with a target recovery of 10,000 nuclei. Library concentration was determined using Qubit (ThermoFisher) and fragment size distributions were assessed using TapeStation (Agilent). Libraries were pooled according to their preparation batches and sequenced on the Illumina NovaSeq X Plus platform using 25B lanes to achieve a target sequencing depth of approximately 10,000 reads per nucleus.

### snRNA-seq data processing and analysis

FASTQ files were processed using DRAGEN Single-Cell RNA v4.4.6000 (Illumina, BaseSpace Sequence Hub), with intron inclusion enabled (--scrna-count-introns=true). This pipeline performs demultiplexing, cell barcode and UMI error correction, sequence alignment to mm10 reference genome, and generation of gene-by-nucleus count matrices. Gene expression matrices were analyzed using Seurat v5 ^82^ in R (v4.5.2). Following quality control, an average of approximately 2,700 high-quality nuclei per sample were retained after quality control. Nuclei with fewer than 200 detected genes, more than 8,000 detected genes, or >1% mitochondrial transcript content were excluded. Each sample was normalized, scaled, and subjected to principal component analysis (PCA) on the top 2,000 highly variable genes, followed by graph-based clustering (resolution = 1.2) and UMAP embedding on the first 15 principal components. Ambient RNA contamination was identified and corrected independently for each sample using scCDC (^83^, restriction factor = 0.4); corrected expression matrices were used for all subsequent analyses. The 24 samples were integrated by canonical correlation analysis (CCA) in Seurat. The integrated data was normalized, scaled, clustered (resolution = 0.8) and visualized by UMAP using the first 30 principal components. Cell identities were assigned by label transfer using stage-matched reference Mouse Retina Cell Atlas ^29^ and confirmed against canonical marker genes for each cell class (rod, cone, bipolar, amacrine, RGC, Müller glia). Lowly-abundant cell types (total nuclei number < 100) and unknown cells (maximum prediction score < 0.5) were excluded for downstream analysis due to insufficient coverage.

### Differential gene expression analysis and rhythm detection

scCDC-corrected counts were aggregated into pseudobulk profiles by biological sample within each cell type. Genes with at least 10 counts in at least six samples were retained for downstream analyses. Unwanted variation was estimated using RUVSeq (RUVs, *k* = 2; ^84^). Differential expression across Zeitgeber time (ZT) was assessed using edgeR ^85^ to identify *rhythmic genes* with FDR < 0.05, and rhythmic parameters such as phase were identified using a modified harmonic regression framework adapted from dryR ^62^. In both analyses, the RUVSeq-derived unwanted variation factors and sex were included as covariates. To analyze the mouse bulk RNAseq retina data from ^24^ using the same pipeline, we downloaded raw data files using the deposited data from the Genome Sequence Archive of the China National Center for Bioinformation using the accession number CRA003588.

### Circadian phase-specific gene ontology enrichment analysis

Gene ontology (GO) biological process enrichment analysis was performed to identify functional pathways associated with genes peaking within specific phase intervals in each retinal cell type. Genes were grouped into six phase bins (ZT0-4, 4-8, 8-12, 12-16, 16-20, and 20-24) using the enrichGO function in the clusterProfiler package ^86^ with the org.Mm.eg.db annotation database. Benjamini-Hochberg correction was applied to account for multiple testing. For visualization, enriched GO terms with fewer than two associated genes were excluded. The three most significantly enriched biological processes within each circadian phase were selected based on adjusted *P*. *value* and displayed as circular heatmaps.

## Author Contributions

MNM and PK contributed to tissue collection. MNM processed the snRNA-seq data samples. GM contributed initial stages of analysis from DRAGEN to differential expression, and MNM contributed to data visualization. SA contributed immunohistochemistry and analysis. MNM, PK, and VMW contributed to the bulk of the writing and editing.

## Acknowledgements

Mice were obtained from The Jackson Laboratory. Retina illustration was designed by Seth D. Lammert.

## Data availability

The data generated in this study are available through a publicly accessible Shiny application at [Mouse retina circadian scRNA-seq gene expression viewer](https://vikkiweake.shinyapps.io/shinyr-1/), which allows users to explore cell type-specific diurnal gene expression patterns in the mouse retina. High-throughput sequencing data is available at the Gene Expression Omnibus (GEO). GSE324665: 4-month-old mouse retina single-nuclei RNA-seq (24 samples).

