## Supplemental Figures for "Defining cell type-specific diurnal gene expression in the mouse retina"

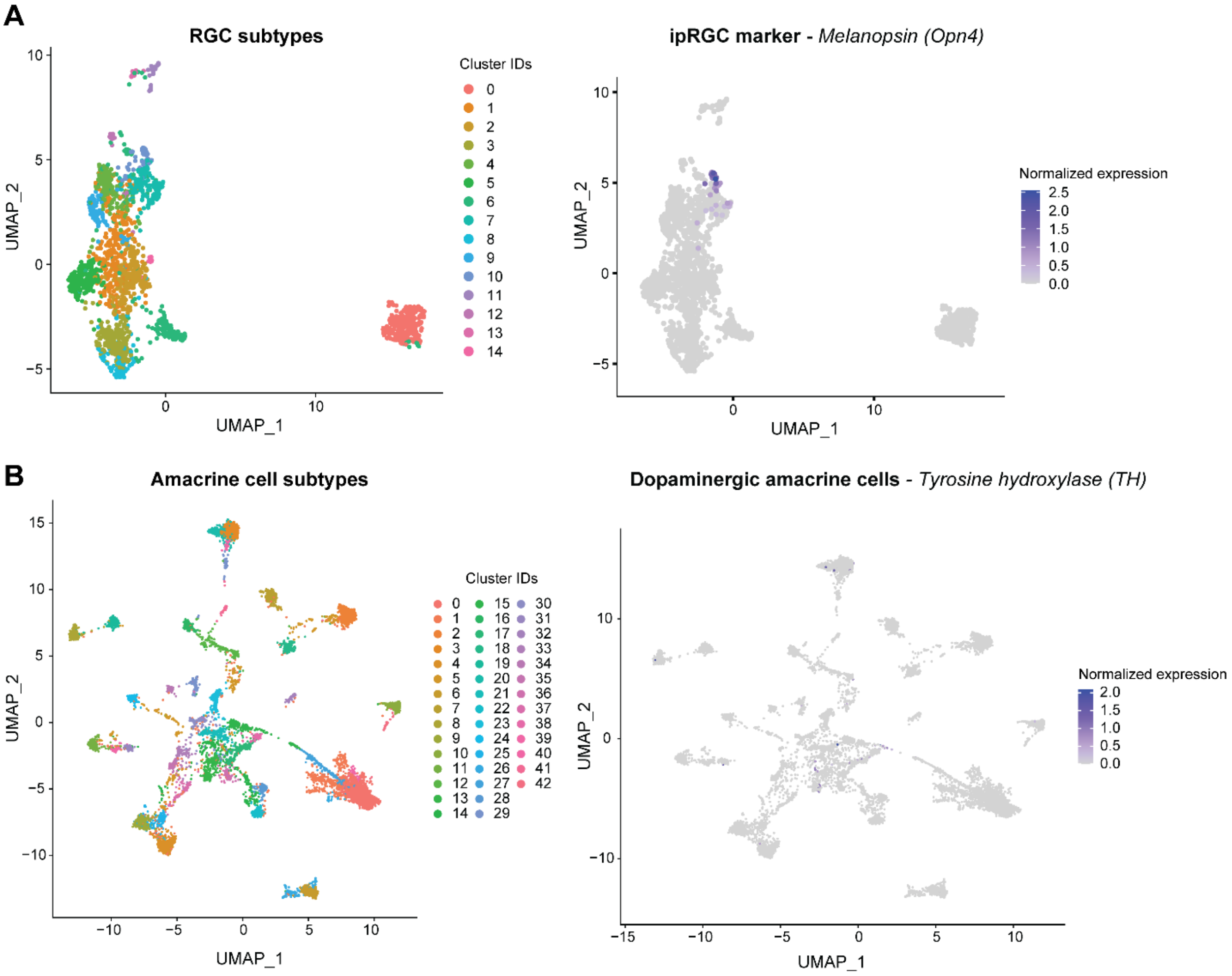


### S1. Annotation of low-abundance retinal neuronal subpopulations

UMAP visualizations of retinal ganglion cell (RGC) and amacrine cell subpopulations. Expression of *melanopsin* (*Opn4), a* marker of ipRGCs, and *tyrosine* hydroxylase (*TH*), a marker of dopaminergic amacrine cells, is shown to support the identification of these specialized subpopulations. These subtypes were detected in the dataset but were not included in downstream cell type-specific gene expression analyses because their low abundance limited the available sequencing coverage.

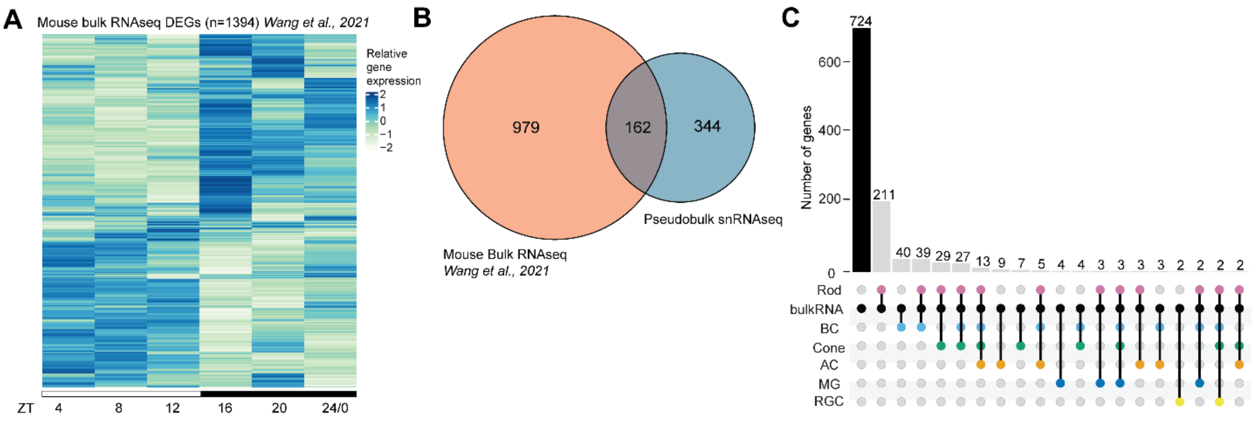


**S2. Comparison of rhythmic gene expression between the present study and published mouse retina transcriptomic data.**

(A) Heatmap showing relative expression of rhythmic genes identified following re-analysis using edgeR (FDR < 0.05) and dryR for amplitude and phase values, of the mouse retina bulk RNA-sequencing data from *Wang et al., 2021*. Expression values are represented as Z-scores calculated by scaling expression for each gene across the six timepoints. Note that collection timepoints are off by two hours from our snRNA-seq data. (B) Venn diagram showing the overlap of rhythmic genes identified in the Wang et al. dataset and our pseudobulk analysis. (C) Upset plot showing the overlap of rhythmic genes identified across individual retinal cell types in the present study and the bulk RNA-sequencing dataset from Wang et al.

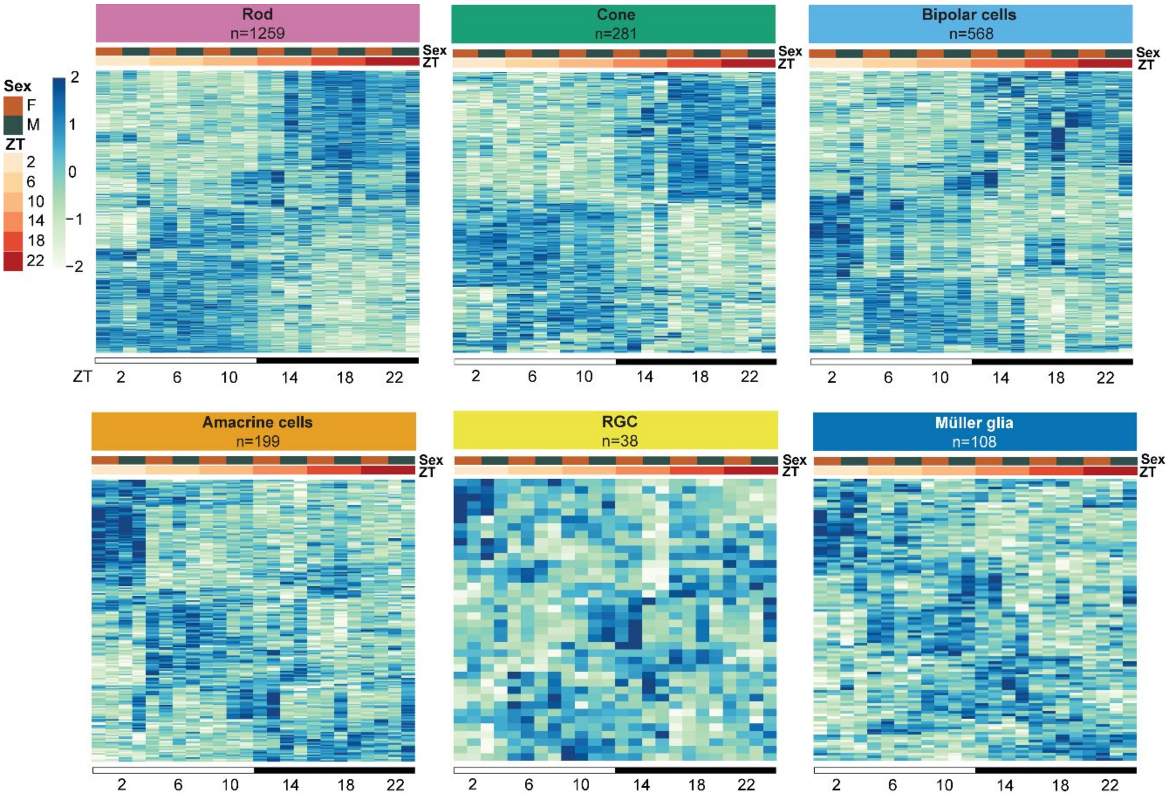


### S3. Replicate-level expression patterns of rhythmic genes across retinal cell types

Heatmaps showing the relative expression of rhythmic genes across the diurnal day in individual male and female biological replicates for each retinal cell types. Expression values are represented as Z-scores calculated by scaling expression values for each gene across the six ZT collection timepoints. Individual replicates are displayed separately to illustrate variability in expression patterns across biological replicates and between sexes.
